# How brain reacts to targeted attack at a hub region

**DOI:** 10.1101/767863

**Authors:** Wenyu Tu, Zilu Ma, Yuncong Ma, Nanyin Zhang

**Author notes:** Address for correspondence: Nanyin Zhang, PhD, Professor of Biomedical Engineering and Electrical Engineering, Lloyd & Dorothy Foehr Huck Chair in Brain Imaging, The Huck Institutes of the Life Sciences The Pennsylvania State University, W-341 Millennium Science Complex, University Park, PA 16802, USA.

## Abstract

The architecture of brain networks has been extensively studied in multiple species. However, exactly how the brain network reconfigures when a local region stops functioning remains elusive. By combining chemogenetics and resting-state functional magnetic resonance imaging (rsfMRI) in awake rodents, we investigated the causal impact of acutely inactivating a hub region (i.e. dorsal anterior cingulate cortex) on brain network properties. We found that disrupting hub activity profoundly changed the function the default-mode network (DMN), and this change was associated with altered DMN-related behavior. Suppressing hub activity also impacted the topological architecture of the whole-brain network in network resilience, segregation and small worldness, but not network integration. This study has established a system that allows for mechanistically dissecting the relationship between local regions and brain network properties. Our data provide direct evidence supporting the hypothesis that acute dysfunction of a brain hub can cause large-scale network changes. This study opens an avenue of manipulating brain networks by controlling hub-node activity.

## Introduction

The mammalian brain is a highly inter-connected network system with distributed brain regions orchestrating to support normal function and mediate complex behavior (*1*). The brain network is organized to maintain an optimal balance of information segregation and integration, which simultaneously maximizes the efficiency of distal communication via long-range connections and minimizes the wiring cost through clustered local processing ((*2, 3*), but also see (*4*)). Such organization is well conserved in multiple species including rodents, primates and humans (*5-7*).

It is becoming increasingly clear that altered brain network organization is tightly linked to brain disorders (*8-10*). However, the neural substrates causing these large-scale network changes remain unknown. A key hypothesis is the dysfunction of brain hub regions. Hubs are brain regions that have high degrees of connections with the rest of the brain (*11, 12*). Because of their central roles, dysfunction of hub nodes can change global integrative process, and has been hypothesized to be a direct cause of altered brain network properties and pathophysiology of brain disorders (*13-15*). For instance, brain network analysis in patients with Alzheimer’s Disease showed that amyloid-beta deposition mainly accumulated in functional hubs (*16*). Therefore, comprehensively understanding the causal relationship between hub activity and brain network organization is critical. However, directly testing this hypothesis in humans is challenging, as selectively altering activity in a hub and dissecting its causal impact on brain networks are difficult.

This obstacle can be overcome in experimental animals using neuroscience tools like Designer Receptors Exclusively Activated by Designer Drugs (DREADDs) and resting state functional magnetic resonance imaging (rsfMRI). DREADDs introduce genetically encoded modified muscarinic G-protein-coupled receptors into living neurons, which allow reversible *in vivo* manipulation of neuronal activity of a defined brain region for hours (*17*). rsfMRI measures brain-wide resting-state functional connectivity (RSFC), and provides comprehensive assessment of brain network properties (*5, 18*). Thus, combining these two techniques offers a system that allows us to characterize the reconfiguration of the brain network as the neural activity of a network node is selectively perturbed. Given the highly conserved brain network architecture in the mammalian brain (*5-7*), the results are translatable to humans.

Here we investigate the causal impact of manipulating a hub region, dorsal anterior cingulate cortex (dACC), on brain network function and organization, as well as behavior using an awake rodent model established in our lab (*19-22*). Imaging awake rodents avoids the confounding effects of anesthesia (*23-26*), and permits linking imaging to behavioral data (*21, 27*). dACC is selected because it is a known functional and anatomical hub in both human and rat brain (*11, 19, 28, 29*). We examine the impact of suppressing the dACC on the default mode network (DMN), which is composed of a group of distributed brain regions exhibiting higher activity at rest (*30, 31*). DMN function is tightly linked to self-referential mental activity in humans, and its abnormality has been linked to numerous brain disorders (*32, 33*). DMN has also been reported in rodents and primates (*6, 34-37*), although our understanding of DMN in animals is mostly limited to its anatomical resemblance to the DMN in humans (*6, 35*). Since dACC is a hub node in the rodent DMN (*34*), manipulating dACC activity provides an avenue for understanding the rodent DMN at the functional level. Our data show that suppressing the dACC disrupts activity and connectivity across the whole DMN, and DMN activity changes are correlated with altered DMN-related behaviors. These data suggest that, like humans, DMN in rodents is a functional network with coordinated activity to mediate behavior. Furthermore, we demonstrate that suppressing the dACC impacts the organization of the whole-brain network including network resilience, segregation and small worldness, but not network integration. All these changes are absent in sham rats or when a non-hub region is suppressed. Taken together, this study provides direct measurement of the brain’s response to targeted attack at a hub region, and presents a comprehensive framework demonstrating the pivotal role of hubs in the brain network.

## Results

Animals were stereotactically injected with adeno-associated viruses (AAVs) expressing inhibitory G-protein coupled hM4Di receptor with a pan-neuronal synapsin promoter (AAV8.hSyn.hM4Di.mCherry, Addgene). After 4-6 weeks of recovery and protein expression (Fig. S1), animals received either Clozapine-N-Oxide (CNO) or saline 30 min before electrophysiology recording, rsfMRI scanning or behavioral tests. Saline and CNO sessions were separated by at least 3 days in a random order. The experimental procedure is summarized in Fig. 1.

**Figure 1.**
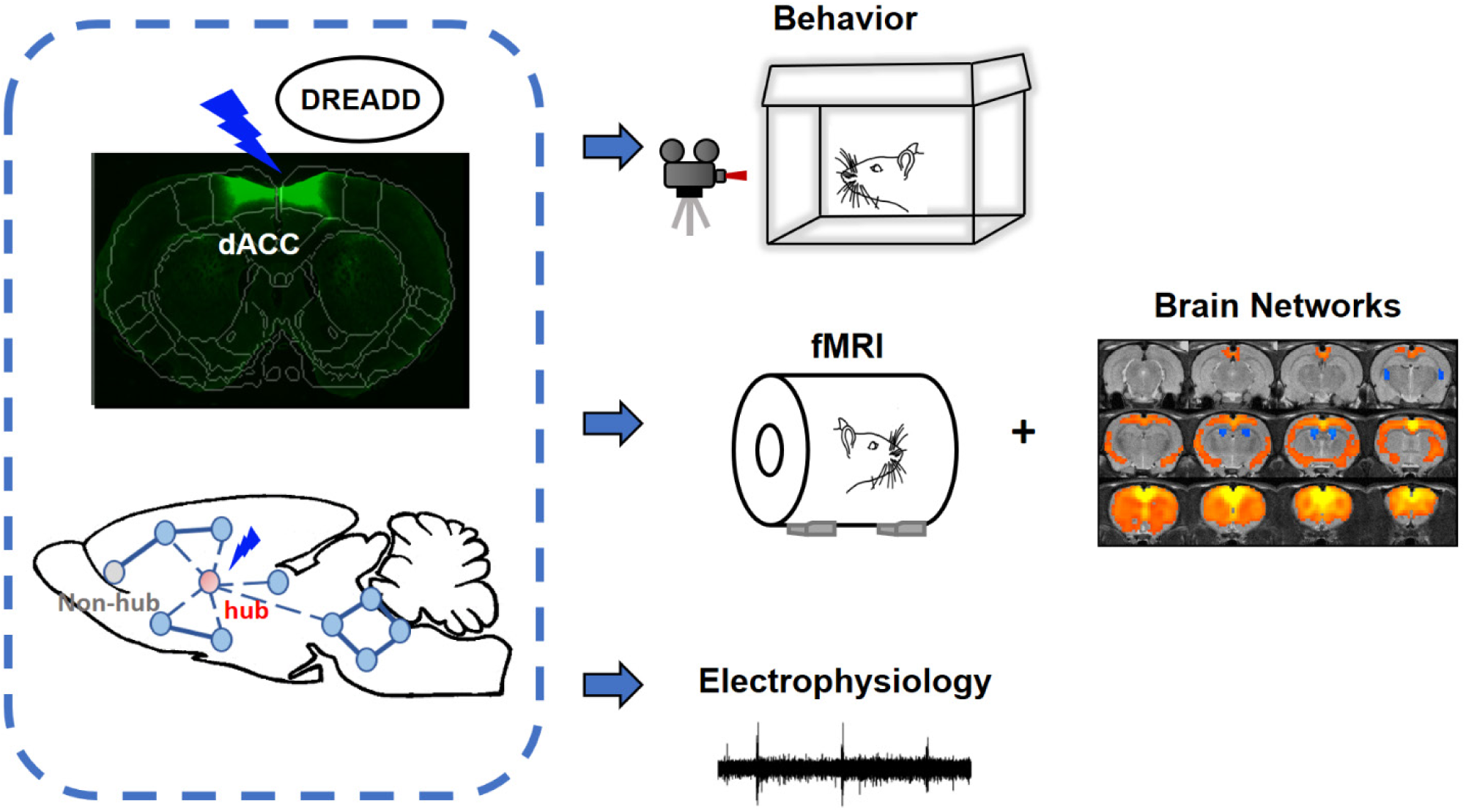
Schematic diagram of the experimental paradigm.

### DREADD suppressed both evoked and spontaneous neural activities in local regions

We first validated the inhibitory effect of the DREADD using a visual stimulation paradigm (Fig.2). AAVs expressing inhibitory DREADD (AAV8.hSyn.hM4Di.mCherry, Addgene) were injected into the superior colliculus (SC). After recovery and DREADD expression, neural activity in the SC was activated by visual stimuli (Figs. 2A-D, 1 flash/trial and 5 flashes/trial, 100ms per flash, 10 sec per trial, 15 trials per animal) and measured by electrophysiology. SC firing rates were significantly reduced 30 min after CNO injection (two-sample tests, 1 flash/trial, t = 10.30, p = 8.8 ×10^−17^, Figs. 2B & 2D; 5 flashes/trial, t = 13.48, p = 4.1×10^−23^, Fig. 2C & 2D). In addition, spontaneous neural activities including spiking activity and local field potential (LFP), quantified by the electrophysiology data 5s before the onset of visual stimulation in each trial, were significantly dampened in CNO-injected rats (MUA: t = 16.15, p = 1.041×10^−36^; LFP: t = 5.84, p = 8.6 × 10^−8^; Figs. 2E, 2F & 2G). The LFP spectrograms after saline and CNO injections in a representative rat were also shown in Fig. S2. These data collectively confirmed the inhibitory effect of the DREADD on evoked and spontaneous neural activities in rats.

**Figure 2.**
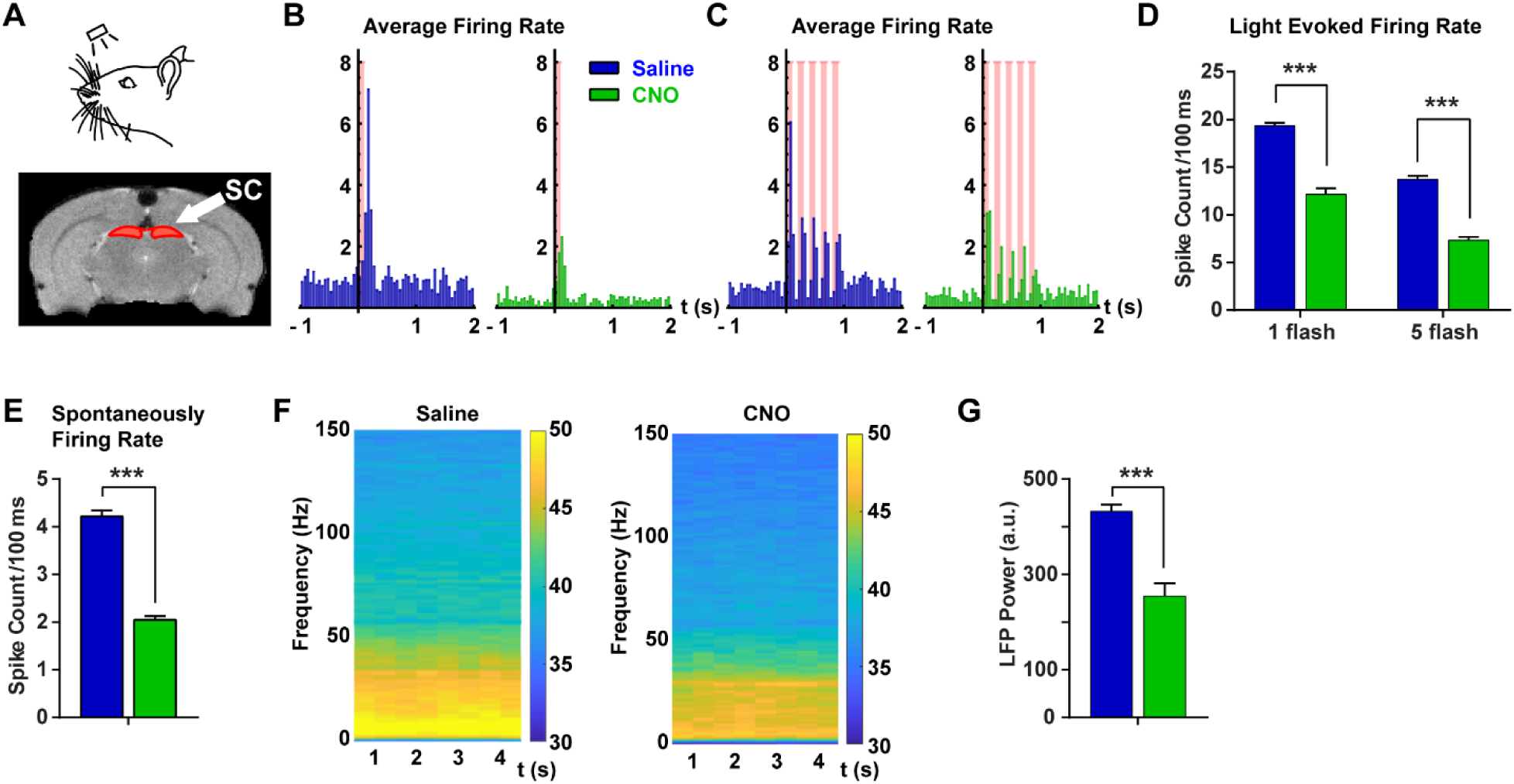
Validation of the inhibitory effect of DREADDs. **A**) Visual stimulation paradigm; Spiking activity in the SC in response to **B**) 1 flash (100ms/flash) and **C**) 5 flashes (100ms/flash) of light stimulation after saline and CNO injections (15 trials per animal, per condition, number of animals = 3); **D**) DREADD suppressed evoked firings (two-sample tests, 1 flash/trial, t = 10.30, p = 8.8 ×10^−17^; 5 flashes/trial, t = 13.48, p = 4.1×10^−23^); **E**) DREADD suppressed spontaneous firings during 5s before visual stimulation (t = 16.15, p = 1.041×10^−36^); **F**) Averaged spectrograms after saline (left) and CNO (right) injections; **G**) Chemogenetic inhibition reduced local field potential power (3-300Hz) in the SC (t = 5.84, p = 8.6 × 10^−8^). ***: p < 0.005.

### Suppressing the dACC reduced its rsfMRI signal, RSFC and local network organization

The bilateral dACC was transfected by inhibitory DREADDs under a pan-neuronal synapsin promoter (AAV8.hSyn.hM4Di.mCherry, Addgene). Sham rats received a control virus (AAV8-hsy-GFP, Addgene). Animals were scanned using rsfMRI twice in the awake state, with each session starting 30 min after either CNO or saline injections. The two imaging sessions were separated by at least 3 days with a randomized order. There was no difference in motion level during rsfMRI scanning between CNO and saline injection conditions in both DREADD and sham groups (Fig. S3).

Fig. 3A shows that DREADD suppression of the dACC reduced its resting-state blood-oxygenation-level dependent (BOLD) amplitude (std normalized by mean, t = 2.00, p = 0.048), again confirming the inhibitory effect of the DREADD on spontaneous neural activity. Suppressing the dACC also affected its functional connectivity, reflected by generally reduced RSFC in dACC seedmaps (for each map: one sample t test, p<0.0005, linear mixed model, false discovery rate (FDR) corrected) after CNO injection relative to saline injection (Fig. 3B). We further examined the impact of suppressing the dACC on its local information processing, assessed by the modular structure, which is defined as a group of nodes with significantly greater within-module connections than between-module connections (*38*). The dACC-related module shrunk in size in DREADD rats after CNO injection relative to saline injection at high graph densities (p < 0.05, linear mixed model, Fig. 3E), suggesting that suppressing the dACC mainly dropped relatively weaker connections in the dACC module.

**Figure 3.**
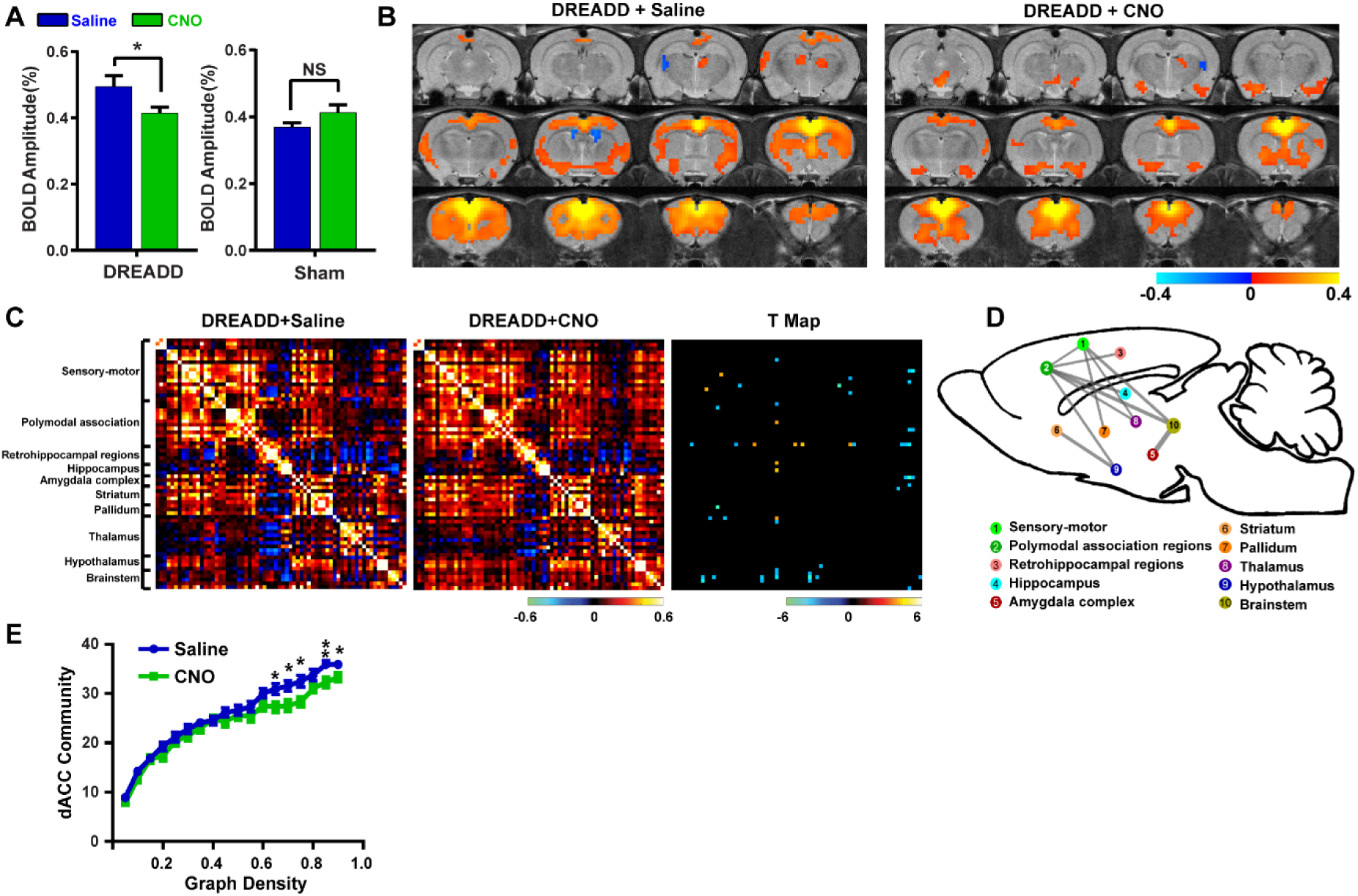
Suppressing dACC reduced its spontaneous BOLD signal, RSFC, and local network structure. **A**) Amplitude of spontaneous BOLD activity in DREADD (19 animals, 78 scans) and sham groups (8 animals, 41scans); **B**) Averaged seedmaps after saline and CNO injections (for each map: one-sample t-test, linear mixed model, p < 0.0005, FDR corrected); **C**) Left: averaged ROI-wise RSFC matrices after saline and CNO injections in the DREADD group. Right: connections exhibiting significant RSFC change (two-sample t-test, linear mixed model, p < 0.05, FDR corrected). **D**) These connections were grouped into multiple brain systems. Line thickness represents the total number of connections with significant RSFC changes between the two systems; **E**) The size of the dACC-associated community structure as a function of graph density in the DREADD group after saline and CNO injections (two-sample t-test, linear mixed model). *: p < 0.05; **: p < 0.01.

In addition to dACC-related circuits, we assessed RSFC between regions of interest (ROIs) across the whole brain. Multiple distributed connections exhibited altered RSFC (two-sample t-tests, p < 0.05, linear mixed model, FDR corrected) with lower RSFC in the majority of these connections (i.e. blue elements in the T map, Figs. 3C & 3D), indicating that the impact of suppressing neural activity in a hub can go beyond the hub itself and propagate to neural circuits across the whole brain. This ripple effect is consistent with the report in human patients with focal brain lesions to critical locations (*39*).

None of these changes (i.e. resting-state BOLD amplitude, dACC-related RSFC and community size) were observed in sham rats when comparing CNO-to saline-injection conditions (Fig. S4), which suggests that CNO had minimal off-target effects in our system, ruling out its potential confounding effects on rsfMRI data (*40*).

### Suppressing dACC altered activity and connectivity in the DMN

Given that dACC is a major hub node in the functional network of DMN, we examined the impact of inactivating the dACC on the DMN organization in awake rodents. We first mapped the DMN using *fractional amplitude of low-frequency fluctuations* (fALFF), defined by voxel-wise low-frequency spectrum power (0.01–0.08 Hz) normalized by the full-spectrum power (*41*) of the rsfMRI signal. This well-established method measures the amplitude of regional spontaneous brain activity. Thus, it can reliably identify brain regions with higher activity at rest, and map the spatial pattern of the DMN (*41*).

Fig. 4 displays the DMN spatial maps in DREADD and sham rats after saline and CNO injections, respectively. Brain regions highlighted in the network included the ACC, posterior cingulate cortex (PCC), prelimbic cortex (PL), retrosplenial cortex (RSC), orbital cortex (ORB), posterior parietal cortex (PPC), as well as subcortical regions of the basal forebrain (BF) and hypothalamus (Hypo). This DMN pattern was reproduced with a completely different analysis method (i.e. independent component analysis, Fig. S5) previously used by other groups, and well agreed with the rodent DMN pattern reported in the literature (*34-36*). Notably, we found that the BF was a prominent node in the rodent DMN, which was not observed in other rodent DMN mapping studies, but was reported in an electrophysiology study that demonstrated that the BF is a key DMN node and critical in regulating DMN-related behaviors in rats (*42*).

**Figure 4.**
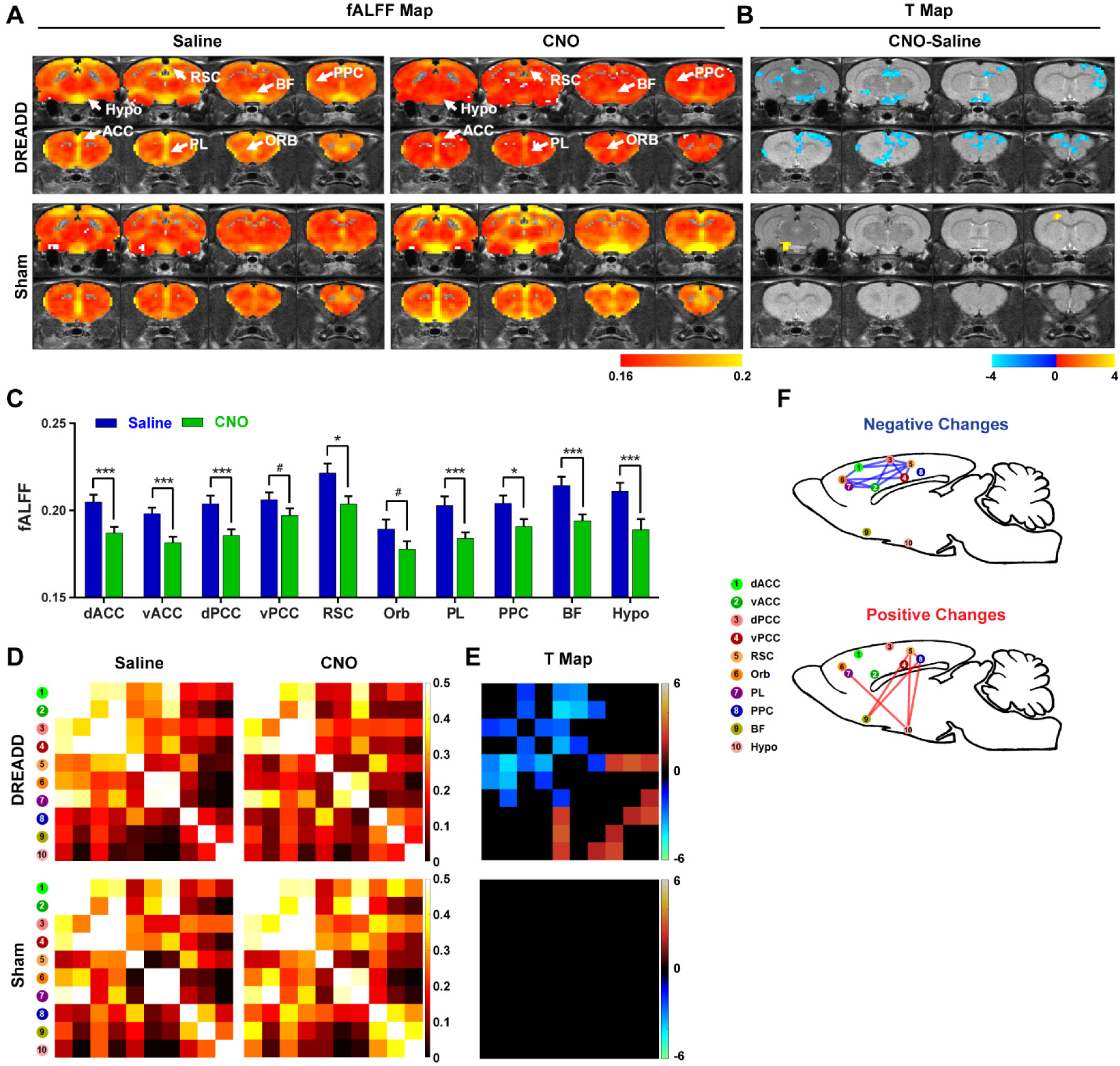
Suppressing dACC disrupted activity across the DMN. **A**) DMN constructed using fractional amplitude of low-frequency fluctuations (fALFF) after saline and CNO injections; **B**) Voxel-wise fALFF changes after DREADD inhibition (p < 0.05, linear mixed model, family-wise error corrected); C) ROI-wise fALFF changes after DREADD inhibition (*: p < 0.05; **: p < 0.01; ***: p < 0.005; #: p = 0.11). **D**) RSFC between DMN regions in the DREADD and sham groups after saline and CNO injections, respectively; **E**) RSFC changes between DMN nodes after dACC inhibition (p < 0.05, linear mixed model, FDR corrected); **F**) Negative and positive RSFC changes after dACC inhibition (p < 0.05, linear mixed model, FDR corrected).

Intriguingly, suppressing one node in the DMN (i.e. dACC) dampened the activity of virtually the entire network (Fig. 4A, top row), reflected by reduced BOLD fALFF in the ACC, dPCC, RSC, PL, PPC, BF and Hypo after CNO injection in DREADD-suppressed rats (Fig. 4B, top row, p < 0.05, Family-wise error corrected, and Fig. 4C). Again, no regions displayed reduced spontaneous activity in sham rats after CNO injection (Figs. 4A & 4B, bottom row). We also calculated RSFC between every pair of nodes in the DMN including the dorsal and ventral ACC, dorsal and ventral PCC, RSC, ORB, PL, PPC, as well as BF and Hypo (*34*) in both DREADD and sham rats after saline and CNO injections, respectively (Fig. 4D). Cortical nodes generally displayed significantly reduced RSFC (p < 0.05, FDR corrected, Figs. 4E & 4F), except that the RSC exhibited increased RSFC with the PPC, BF and Hypo after CNO injection. In contrast, both subcortical nodes (BF and Hypo) showed increased RSFC after dACC suppression (Figs. 4E & 4F). Taken together, these data revealed that the DMN was significantly reorganized when dACC activity was suppressed, suggesting that the rodent DMN is a functional network with coordinated activity, and the dACC is a pivotal node in this network.

### Suppressing dACC altered DMN-related behaviors in animals

Considering that suppressing dACC significantly altered DMN activity in awake rodents, we hypothesize that it also changes DMN-related behavior, measured by quiet restfulness in rodents. Nair and colleagues demonstrated that quiet restfulness in home-cage is characteristic behavior to DMN activity, evidenced by elevated neural activity in the DMN, including ACC and BF, during this behavioral state (*42*). Robust DMN activation during quiet restfulness was also reported in chimpanzees (*43*). Therefore, we evaluated quiet restfulness, defined by continuous immobility for at least 2s, in our animals. After either CNO or saline injection, the rat was put in the home cage for 45min and was video-recorded. The animal’s behavior was analyzed using behavioral tracking software (ANY-maze, Stoelting Co., Wood Dale, IL). After CNO injection, quiet restfulness was significantly reduced in the rat (Fig. 5), reflected by significantly lower immobile time for at least 2s (t = 3.991, p = 0.0009), but higher distance travelled (t = 4.294, p = 0.0004), mean speed (t = 4.811, p = 0.0001) and total mobile time (t = 3.991, p = 0.0009). Rats also displayed significantly higher rearing after CNO injection (t = 2.80, p = 0.01), likely reflecting an increase in vigilance and/or exploratory behavior. Reduced quiet restfulness remained consistent during the last 15 min of testing, suggesting that these behavioral changes were not due to the environmental changes at the beginning of the test (Fig. S6).

We also examined whether DMN activity changes could explain altered DMN-related behaviors. In 8 animals, both behavioral and imaging data were collected after DREADD expression. Notably, both data were collected in the awake state, which permits linking results of these two measures. We found that fALFF changes in the dPCC and PL were significantly correlated to changes in all measures of DMN-related behavior including distance travelled (dPCC: r = 0.80, p = 0.016; PL: r = 0.79, p = 0.019), mean speed (dPCC: r = 0.85, p = 0.0077; PL: r = 0.84, p = 0.0095), mobile time (dPCC: r = 0.74, p = 0.036; PL: r = 0.80, p = 0.017) and immobile time (dPCC: r = 0.74, p = 0.036; PL: r = 0.80, p = 0.017). These data collectively demonstrate that dACC suppression alters DMN activity, and DMN activity changes can predict alteration in DMN-related behaviors.

**Figure 5.**
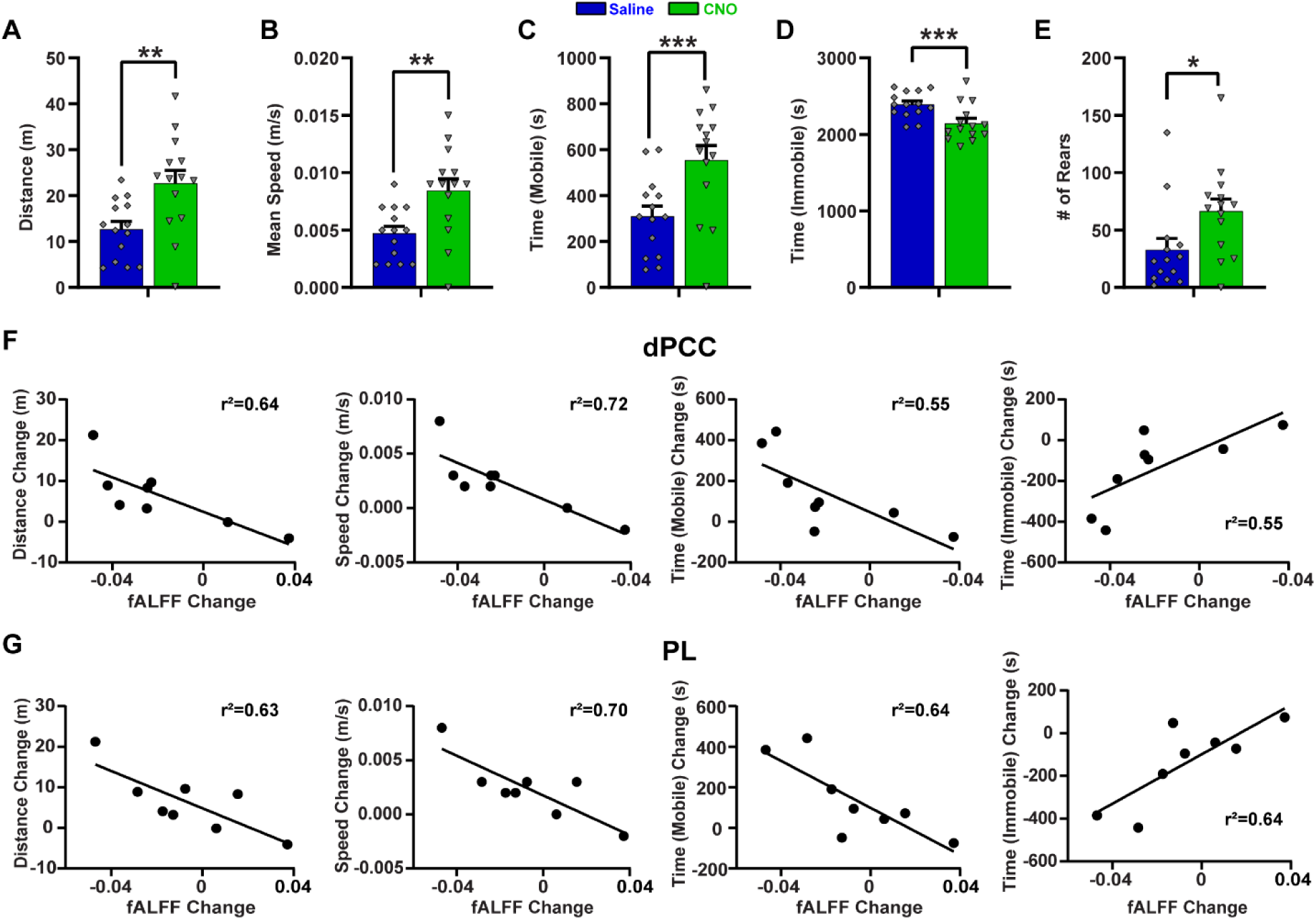
Suppressing dACC changed DMN-related behavior (n=14). **A**) Total distance traveled in homecage (45 min); **B**) Mean speed; **C**) Total mobile time; **D**) Total immobile time for at least 2s. **E**) Total rearing behavior. *: p < 0.05; ***: p < 0.005. **F**) Correlations of dPCC activity and DMN-related behaviors. **G**) Correlations of PL activity and DMN-related behaviors.

### Suppressing a hub node changed whole-brain network organization

We further investigated the effect of suppressing the dACC on the organization of the whole-brain network architecture in four aspects: network resilience, functional segregation, functional integration, and small-worldness. The rat brain was parcellated into 68 bilateral ROIs based on the Swanson atlas (*44*). The whole-brain network was constructed by calculating RSFC between every two ROIs, generating a RSFC matrix for each animal at each condition. This RSFC matrix was then binarized based on graph density. Graph topological parameters including assortativity, modularity, global efficiency, and small-worldness were respectively calculated as a function of graph density.

Assortativity measures network resilience. Brain networks with high assortativity are more resistant to local attack such as lesion and neurological degeneration (*12, 45*). Fig. 6 shows that suppressing the dACC considerably reduced the network assortativity, and as a consequence, the whole-brain network became more vulnerable (Fig. 6A). Functional segregation, quantified by modularity, measures the network’s ability of specialized processing. We observed that dACC suppression decreased the modularity of the global brain network, indicating reduced functional segregation (Fig. 6C). Global efficiency measures functional integration, which assesses the ability to integrate specialized information from distributed brain regions. Interestingly, suppressing the dACC did not seem to affect the global efficiency (Fig. 6D). Small-worldness reflects the ability to balance functional segregation and integration. Suppressing the dACC reduced small-worldness of the whole-brain network (Fig. 6B). In summary, we observed that suppression of the dACC affected global brain topology including network resilience, network segregation and small-worldness, but not network integration.

**Figure 6.**
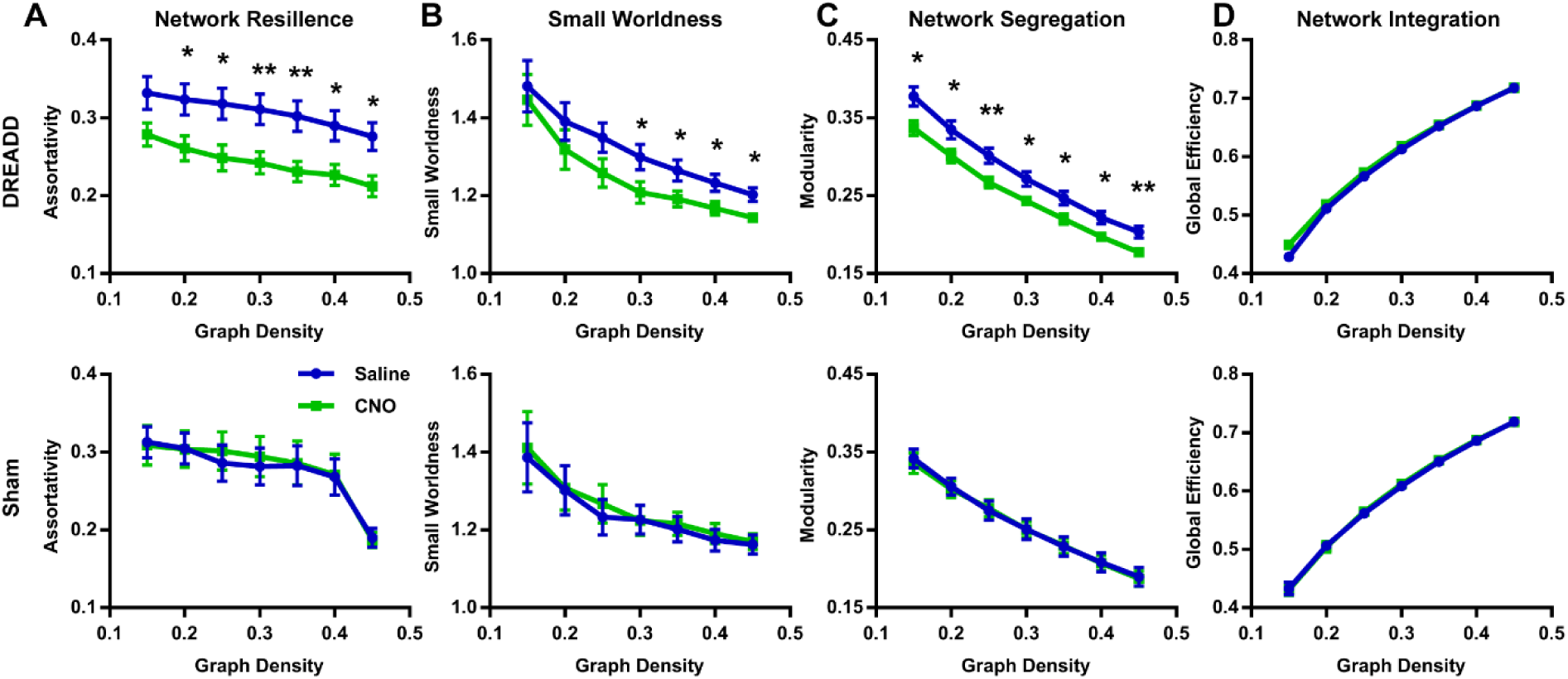
Suppressing a hub node changed whole-brain network topology. **A**) Network resilience; **B**) Small-worldness; **C**) Network segregation; **D**) Network integration. Linear mixed model; DREADD: n = 19, 78 scans; sham: n = 8, 41 scans. *: p < 0.05; **: p < 0.01.

As expected, DREADD suppression of the dACC significantly lowered its degree (Fig. 7), suggesting that dACC lost its hubness in the network. Interestingly, the degree of the ventral RSC was significantly increased after dACC suppression, which is consistent with our data that some RSC connections increased their RSFC within the DMN (Figs. 4E & 4F). Again, no changes in the degree of dACC or vRSC were observed in sham rats (Fig. S7). These results indicate that a new hub can emerge when an existing hub stops functioning, likely due to the compensatory mechanism.

**Figure 7.**
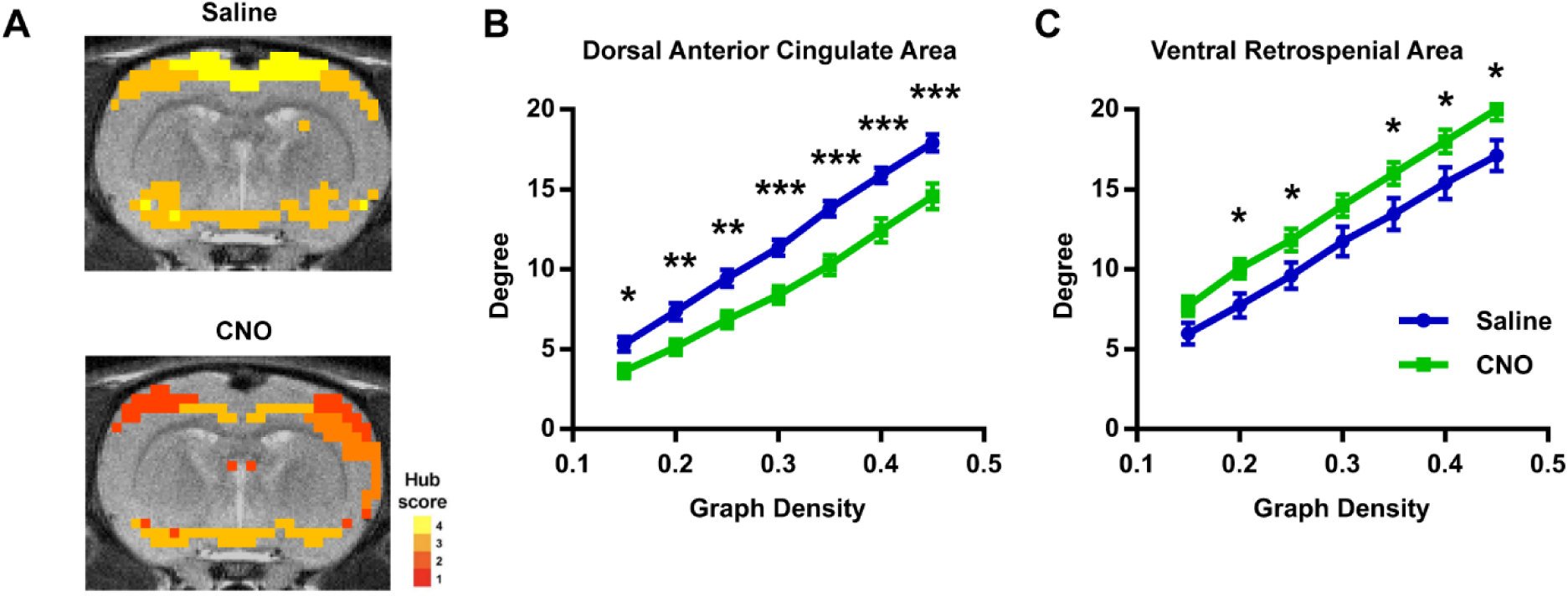
New hub emerged after the inactivation of an existing hub. **A**) Hub scores without and with dACC inactivation, respectively; **B**) The degree of dACC and ventral RSC without and with dACC inactivation (linear mixed model; n = 19, 78 scans). *: p < 0.05; **: p < 0.01; ***: p < 0.005.

### Suppression of a non-hub region did not change whole-brain network properties

To determine whether aforementioned network changes depended on the specific role of the node (i.e. hub versus non-hub), we also suppressed the activity in a non-hub region—primary visual cortex (V1, bilateral) (*19*) using the same pan-neuronal inhibitory DREADDs. After CNO injection, significantly reduced BOLD amplitude in V1 was observed (Fig. 8A), which confirmed the inhibitory effect of DREADD in V1. However, suppression of V1 did not affect fALFF in the DMN (Fig. S8), nor did it cause any whole-brain topological changes including network resilience, functional segregation, functional integration, or small-worldness (Figs. 8B-8E). Taken together, these data suggest that attack on separate brain regions can have differential impact on brain network properties, and brain networks are particularly vulnerable to targeted attack at hub nodes.

**Figure 8.**
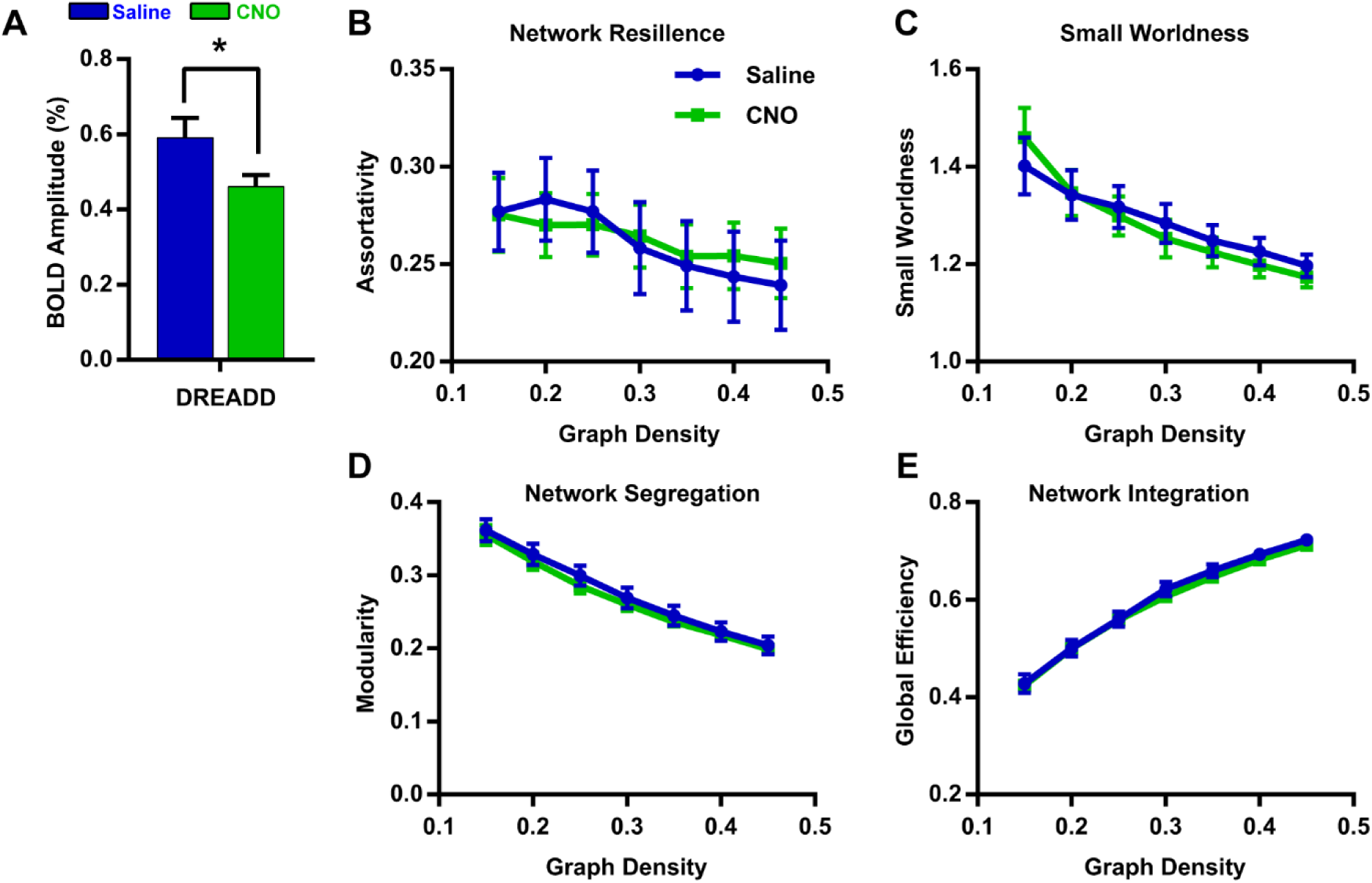
Suppressing a non-hub node did not change whole-brain network topology. **A**) Amplitude of spontaneous BOLD activity in the primary visual cortex after CNO and saline injections (p < 0.05, n=8, 66 scans); **B**) Network resilience; **C**) Small-worldness; **D**) Network segregation; **E**) Network integration.

## Discussion

For over a century, the neuroscience field has been tremendously moved forward by the effort of examining the relationship between damage of specific brain regions and loss of certain brain function. Classical examples include the discovery of the inferior frontal gyrus in speech production by Paul Broca, and the left posterior temporal cortex in language comprehension by Carl Wernicke. Despite this vital advancement, converging evidence suggests that a large number of complex tasks involve numerous cognitive processes and require integrated activity from multiple brain regions (*46*). Therefore, brain function should be viewed as a network phenomenon.

By combining DREADDs and rsfMRI in an awake rodent model, we have established a system that eables us to manipulate the activity in any brain region, and measure the corresponding changes in brain-wide networks. This system allows for mechanistically dissecting the causal relationship between a (or multiple) local brain region(s) and brain network properties. In this study we specifically investigated the role of a hub region (i.e. dACC) in brain networks. We found that disrupting hub activity profoundly changed the function of the DMN (Figs 4), and this change was associated with altered DMN-related behavior (Fig. 5). It also impacted the topological architecture of the whole-brain network in terms of network resilience, segregation and small worldness, but not network integration (Fig. 6). These data offer direct evidence supporting the hypothesis that acute dysfunction of a brain hub can cause large-scale network and behavioral changes. They also provide a comprehensive framework documenting the differential impact of inactivating a hub versus a non-hub node on network dynamics.

### Tight relationship between local regions and brain networks

The activity in local brain regions can be manipulated using tools like chemogenetic, optogenetic, pharmacological and lesion methods. Such perturbations are often used to study the role of a specific brain region in behavior (*47*). Meanwhile, the capacity of manipulating local brain regions can be tremendously expanded when combining with neuroimaging approaches with a global field of view (e.g. fMRI), as the causal impact of local region dysfunction on the rest of the interconnected network can be monitored across the whole brain (*48*). For instance, using DREADDs and rsfMRI, Grayson and colleagues showed that inactivating the amygdala disrupted communication throughout the cortex in anesthetized monkeys (*49*). This observation echoes human patient studies in which lesions of critical focal brain locations caused widespread disruption of the brain modular structure (*39, 50*). These results collectively suggest tight linkage between regional activity and large-scale network properties, and highlight the importance of combining local perturbations and global network measurement. Uniquely, in our study we specifically targeted a hub region to directly test the hypothesis that dysfunction of hub nodes is a direct cause of altered brain network properties (*13-15*). In addition, all data were collected in awake animals, which avoids the confounding effects of anesthesia on brain network function, and permits direct association of network features and behavioral changes. This nature also makes the findings translatable to humans, considering numerous reports of large-scale brain network changes in human brain disorders.

### DREADDs vs lesion

There are several advantages choosing the DREADD over the lesion method for our research purpose. First, DREADDs allow us to specifically and reversibly inhibit a brain region with minimal damage. On the other hand, long-term damage in brain tissue can decrease metabolism and alter local vascular structure. In addition, there can be excessive blood flow in surrounding regions (*51*). All these factors will impact fMRI data measurement and can confound RSFC results. Long-term lesion studies can also be compounded by post-lesion neural plasticity changes resulting from the compensatory mechanism (i.e. secondary effects). Furthermore, most psychiatric disorders are linked to loss of function without tissue damage, making the DREADD method a better translatable model.

### DMN in rodents – beyond anatomical resemblance with other species

We observed significant reconfiguration of the DMN after transiently inactivating one of its hub nodes (i.e. dACC) in awake rodents. DMN activity changes were also linked to altered DMN-related behaviors after dACC suppression. Although the DMN has been extensively studied in humans including its network structure, function and clinical implications, our understanding of the DMN in rodents is limited. The discovery of rodent DMN was mainly based on the anatomical resemblance of the network structure with the DMN in humans and primates (*35, 36*), but its functional role in behavior remains unclear. Nair and colleagues showed that the rat BF exhibited pronounced gamma oscillations during DMN-related behaviors and this activity was suppressed during active exploration of an unfamiliar environment (*42*), suggesting that the BF might be a key DMN node that regulates DMN activity. They further demonstrated that the BF controls DMN-related behaviors by affecting neural activity in the ACC. These results are highly consistent with our findings. First, we also observed prominent involvement of the BF in the rat DMN at baseline, which was missing in previous rodent DMN mapping studies (*34, 35*). Notably, the BF has strong connections with the cortex and subcortical regions like Hypo, and these connected regions substantially overlapped with the DMN revealed in our study (*52, 53*). Second, suppressing the dACC considerably dampened the DMN activity. This result is consistent with the report that DMN hubs are mostly fragile (*54*). Third, reduced DMN-related behaviors resulting from dACC suppression were correlated to activity changes in DMN nodes. Taken together, these results support the concept that, like humans and primates, the DMN in rodents is a functional network with coordinated neural activity from distributed brain regions, and this network might support behavior related to internally oriented brain states.

### Hubs and brain network topology

The topological changes resulting from hub inactivation, including reduced network resilience, segregation and small worldness, are important for our understanding of the role of individual regions in the global brain network organization. A brain network can be modeled as a graph composed of spatially distributed neuronal components as nodes and anatomical or functional associations between components as edges (*2, 3*). Topological features of the brain graph are critical in understanding inter-areal information exchange in health and disease. Although the architecture of brain networks has been extensively studied in multiple species (*5, 55-57*), it remains elusive whether different types of nodes, such as hubs versus non-hubs, play distinct or similar roles in the topology of the global brain network. Our data revealed that changes in a hub, but not a non-hub, can significantly alter brain network topology. These results provide critical insight into understanding the pathogenesis of neurological and psychiatric disorders, which indicate that altered brain topological properties reported in these brain disorders might start from dysfunction of certain hub nodes (*58*). Interestingly, our data showed that network integration was intact after dACC inactivation, likely due to the compensatory effect of the so-called ‘rich club organization’ in the rat brain (*19*). Another possible explanation is that although the dACC has high centrality, it functions as a provincial hub as opposed to a connector hub (*59*). These two types of hubs were demonstrated to have differential impact on brain network properties (*39*), with connector hubs more affecting network integration, and provincial hubs more affecting network segregation. Our result is consistent with the report that the ACC belongs to the “segregation effect” class, as reported in human lesion studies (*60*). Our data also support the notion that neural network communication does not necessarily involve the shortest paths between nodes (e.g. navigation routing) (*61*).

## Methods and Materials

### Animals

44 adult male Long-Evans rats (300-500g) were used in the present study. Animals were housed in Plexiglas cages. Food and water were provided *ad libitum*. The housing room was maintained at an ambient temperature (22-24 °C) with a 12h light:12h dark cycle. All experiments were approved by the Pennsylvania State University Institutional Animal Care and Use Committee.

### Surgery

Aseptic stereotaxic surgery was conducted to infuse the virus to the brain. Rats were anesthetized by intramuscular (IM) injections of ketamine (40 mg/kg) and xylazine (12 mg/kg). Dexamethasone (0.5 mg/kg) and Baytril (2.5 mg/kg) were injected SQ to prevent tissue inflammation and bacterial infections. During the surgery, respiration was controlled by a ventilator with artificial oxygen. The heart rate was monitored by a pulse oximetry (MouseSTAT® Jr., Kent Scientific Corporation), and the body temperature was maintained at 37 °C with a warming pad (PhysioSuite, Kent Scientific Corporation). The DREADD virus (AAV8-hSyn-hM4Di-mCherry, 1 µL at titer ≥ 3×10^12^ vg/mL, Addgene, Watertown, MA) was bilaterally injected into the dACC (coordinates: AP +2, ML +/-0.5, DV −1) and the primary visual cortex (coordinates: AP −6.5, ML +/−3.5, DV −1) for the hub (n = 25) and non-hub (n = 8) groups, respectively ^24^. Sham rats were injected with a control virus (AAV8-hsy-GFP, 1 µL at titer ≥ 3×10^12^ vg/mL, Addgene, Watertown, MA, n = 8)^24^ into the dACC. In addition, separate animals used in the visual stimulation experiment were unilaterally injected with AAV8-hSyn-hM4Di-mCherry (1 µL at titer ≥ 3×10^12^ vg/mL, Addgene, Watertown, MA, n = 3) in the SC (coordinates: AP −7, ML +1.5, DV −3). Rats were allowed to recover for at least four weeks.

### fMRI experiments

Before imaging, animals were acclimated to the MRI environment for 7 days in order to minimize stress and motion during imaging. To better adapt the animal to the restrainer and scanning environment, the acclimation period was gradually increased from 15 min on Day 1 to 30 min on Day 2, 45 min on Day 3, and maintained at 60 min/day for Days 4-7. Detailed procedures for acclimation can be found from our previous publications (*62, 63*). A similar approach to imaging awake animals has also been adopted by other research groups in multiple species (*64-66*). CNO (1mg/kg in saline, dissolved in DMSO, Sigma-Aldrich, St. Louis, MO) or saline (with DMSO) was IP injected 30 min before rsfMRI data acquisition, at a random order with at least three days apart. All imaging sections were conducted at the High Field MRI Facility at the Pennsylvania State University on a 7T Bruker 70/30 BioSpec running ParaVision 6.0.1 (Bruker, Billerica, MA). rsfMRI data were acquired at the awake state using T2*-weighted gradient-echo rsfMRI images using the echo-planar-imaging (EPI) sequence with the following parameters: repetition time = 1000ms; echo time = 15ms; matrix size = 64×64; field of view = 3.2 × 3.2 cm^2^; slice number = 20; slice thickness = 1mm; flip angle = 60; 600 volume each run. Three runs were acquired each rsfMRI session. Raw EPI images were shown in Fig. S9.

### Data analysis

All acquired image data were aligned to a defined atlas using Medical Image Visualization and Analysis software (MIVA, http://ccni.wpi.edu/), and then subjected to motion correlation (SPM12), spatial smoothing (Gaussian kernel, full-width at half-maximum = 1 mm), voxel-wise nuisance regression of motion parameters, as well as the signals from the white matter and ventricles, and bandpass filtering (0.01-0.1Hz). Before these steps, rsfMRI volumes with relative framewise displacement (FD) > 0.25 mm and the preceding and following volumes were removed. In addition, the first 10 volumes of each rsfMRI scan were discarded to ensure steady state of magnetization. rsfMRI scans with > 15% volumes discarded were removed from further analysis.

fALFF was determined by calculating the ratio of the power in the low-frequency range (0.01–0.08 Hz) to that of the full spectrum (0–0.25 Hz). For each voxel, the time course was transformed to the frequency domain using FFT. We calculated the ratio between the square root of power in the range of 0.01–0.08 Hz and that across the entire frequency range. fALFF changes after DREADD inhibition were obtained using voxel-wise two-sample t-tests with family wise error correction (P<0.05, FWE corrected) between data acquired after CNO injection versus saline injection.

Seed RSFC maps were generated using seed-based correlation analysis. Pearson cross-correlation coefficients between the regionally averaged time course from all voxels inside the seed and time courses of individual voxels were calculated. The corresponding correlation coefficients were Fisher transformed, averaged across all scans and animals in each group, and transformed back to r values.

The whole-brain RSFC network was constructed by parcellating the rat brain into 68 anatomical ROIs based on the Swanson Atlas (*44*), which were also grouped into 10 functional systems including amygdala complex, striatum, hypothalamus, hippocampus, thalamus, sensory-motor cortex, polymodal association cortex, brainstem, pallidum, and retro-hippocampal regions. RSFC between each pair of ROIs was calculated by Pearson correlation of regionally averaged time series of the ROIs.

A hub score of each node ranged from 0-4 was determined by the number of the following four criteria the node met: (1) 20% highest degree; (2) 20% highest betweenness centrality; (3) 20% lowest characteristic path length; (4) 20% lowest local clustering coefficient.

Topological parameters of the whole-brain network including the modularity, global efficiency, small-worldness and assortativity were calculated on binarized network across the graph density ranging from 0.15 to 0.45 with a step of 0.05 using the brain connectivity toolbox (BCT) ^30^ based on the following equations:

### Assortativity (45)

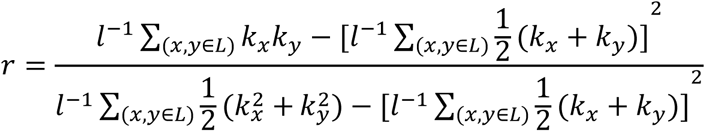

where k_x,_ k_y_ is the degree of node x and y on two opposite ends of an edge; L is the set of all edges within the network, and l is the total number of edges.

### Modularity (67)

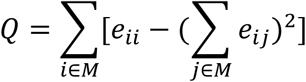

where M represents a set of non-overlapping modules, and e_ij_ is the proportion of between-modules connections of module i and j; e_ii_ is the proportion of within-modules connections of module i.

### Global efficiency (68)

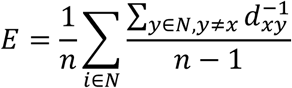

where n is the total number of nodes; N is the set of all nodes. d_xy_ is the shortest path length between nodes x and y.

### Small-worldness (69)

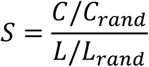

where C and C_rand_ are the clustering coefficients of the testing network and a random network, respectively; L and L_rand_ are the characteristic path lengths of the testing network and a random network.

At each graph density, all topological parameters were determined for each scan. Changes in each parameter was determined by two-sample t-tests (linear mixed model) between the CNO injection and saline injection conditions.

### Behavioral Test

Animals underwent two behavioral sessions with either CNO or saline injection with a random order separated by at least 7 days. Each session was composed of a 45 min home-cage test. Before the test, the animal was acclimated to the behavioral room for 15 min, followed by either CNO (1mg/kg) or saline injection. The animal was then put back to its home-cage in the behavioral room for 45 min. Behaviors including the total distance traveled, mean speed, time of mobility and time of immobility for at least 2s were recorded by an infrared camera and analyzed by behavioral tracking software (ANY-maze, Stoelting Co., Wood Dale, IL).

### Electrophysiology

Electrophysiology recordings were conducted in animals with inhibitory DREADD expressed in the SC. Rats were initially anesthetized by IM injections of ketamine (40 mg/kg) and xylazine (12 mg/kg). An electrode (Neuronexus, Ann Arbor, MI) was then inserted into the SC to measure visually-evoked neuronal response to light stimulation produced by a blue laser source (100 mW, 473 nm, Opto Engine) coupled with an optic fiber, which was placed 5 cm away from the contralateral eye. Before recording, rats were injected with either saline or CNO in each session. During recording, light anesthesia was maintained using isoflurane (∼ 0.75%). Each session included 15 trials of visual stimulation, presented as a single light flash (100 ms per flash) or five flashes (100 ms each flash, 100 ms inter-flash interval) every 10 s. Light stimulation was controlled by a custom Labview program. The electrophysiological signal was amplified and sampled at 25 kHz using a Neuronexus recording system (Neuronexus, Ann Arbor, MI). Raw data were bandpass filtered (MUA: 300-3000 Hz, LFP: 3-300 Hz). Spikes larger than three times of the standard deviation of electrophysiological signal were detected and clustered at a bin size of 50 ms in peristimulus-timed histograms (PSTHs).

### Histology

After electrophysiology, animal was perfused with saline followed by 4% PFA solution. The brain was taken out carefully and stored in solution with 4% paraformaldehyde and 20% sucrose. After fixation, the brain was sliced into 60μm slices. Fluorescent expression in the injection site was examined under microscopy.

## Acknowledgments

We would like to thank Yikang Liu and David Dopfel for their technical support. The present study was partially supported by National Institute of Neurological Disorders and Stroke Grant R01NS085200 (PI: Nanyin Zhang, PhD) and National Institute of Mental Health Grant R01MH098003 and RF1MH114224 (PI: Nanyin Zhang, PhD).

## Author Contributions

N.Z. designed the research; W.T., Z.M. and Y.M. performed animal experiments; W.T. and Y.M. performed data analysis; N.Z. and W.T. wrote the paper.

## Data availability

Data supporting the findings of this manuscript are available from the corresponding author upon reasonable request.

## Supplementary Information

**Figure S1.**
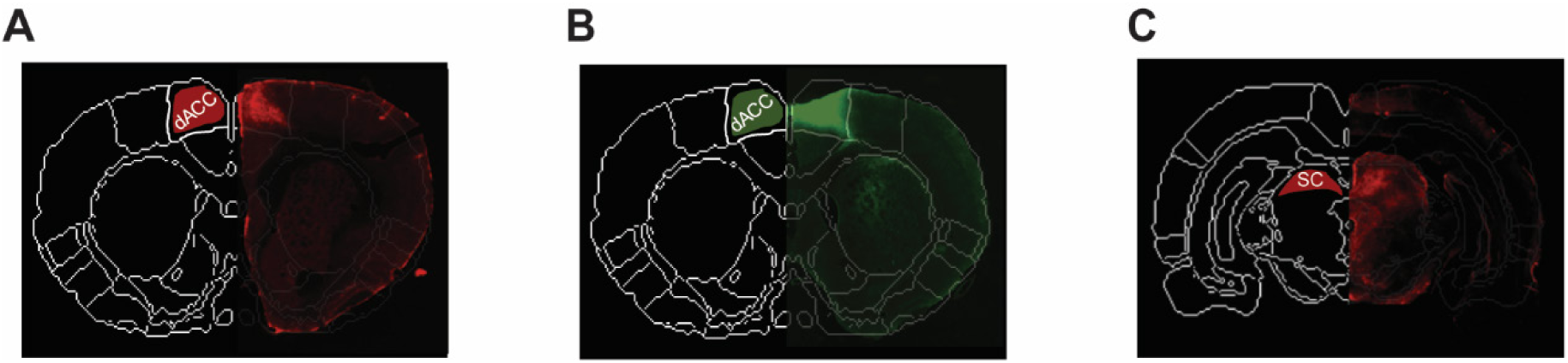
Representative Histology. Protein expression is shown in the right hemisphere. **A**) DREADD expression in the dorsal anterior cingulate cortex (dACC); **B**) GFP in the dACC (i.e. sham rats); **C**) DREADD in the superior colliculus. Anatomical definition is shown in the left hemisphere.

**Figure S2.**
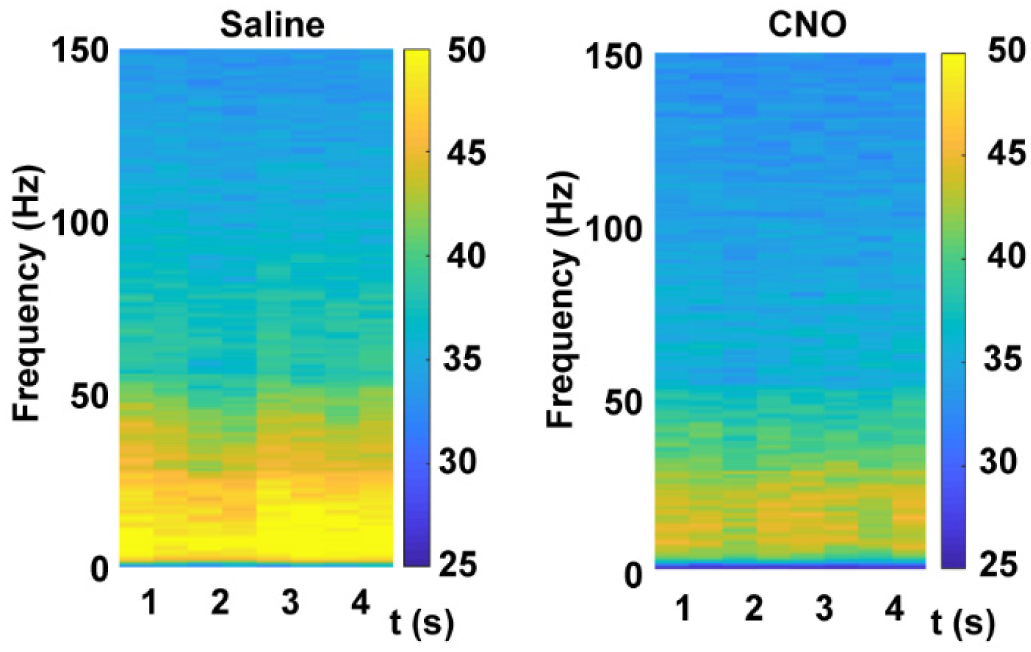
LFP spectrograms after saline and CNO injections in a representative rat with DREADDs infused in the SC.

**Figure S3.**
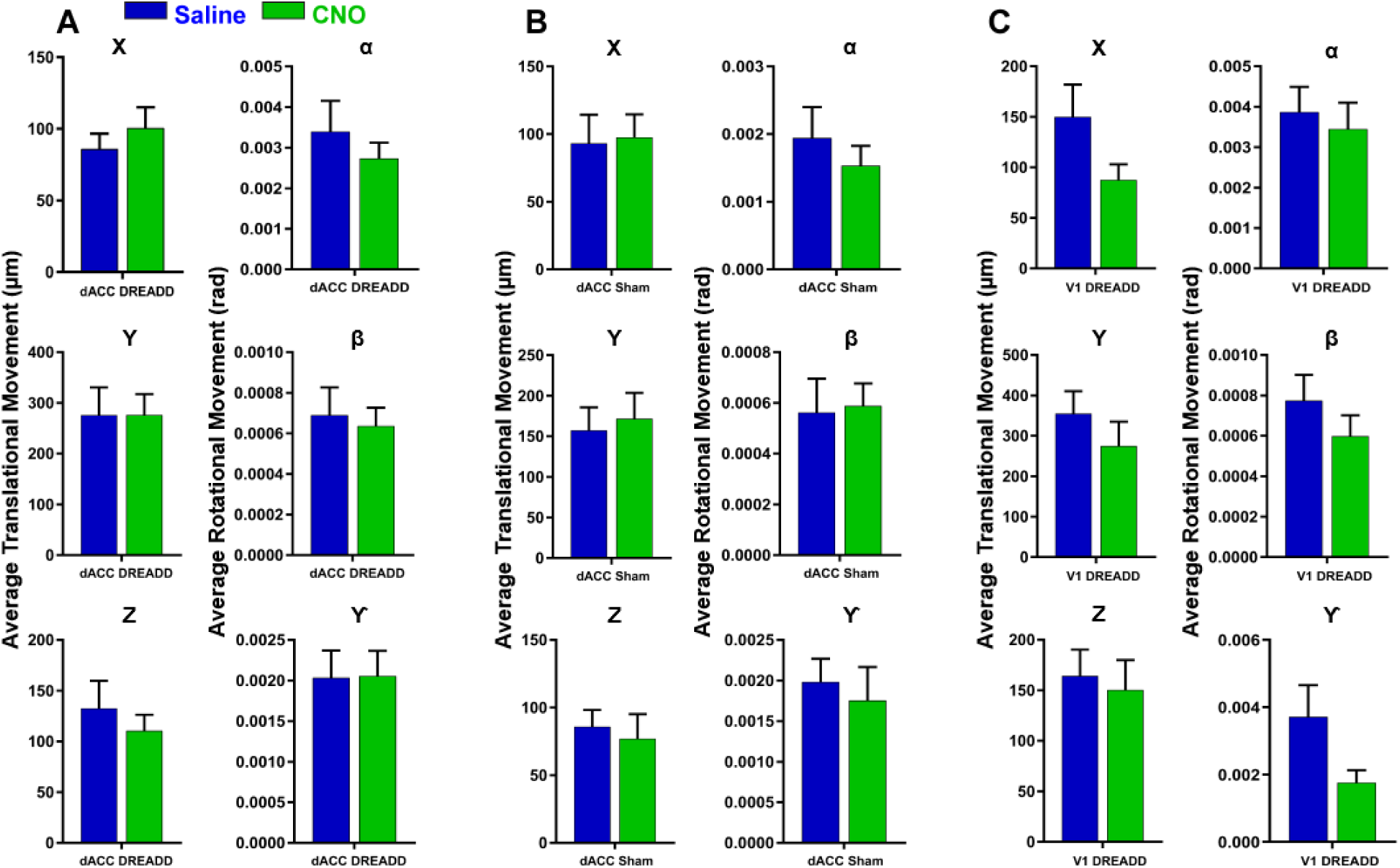
Motion levels in all groups. **A**. dACC DREADD (n=19, 78 scans); **B**. dACC sham (n=8, 41 scans); **C**. V1 DREADD (n=8, 66 scans). Left column: translational movement (in μm). Right column: rotational movement (in rad) (linear mixed model, p > 0.05 for all 6 parameters in all groups).

**Figure S4.**
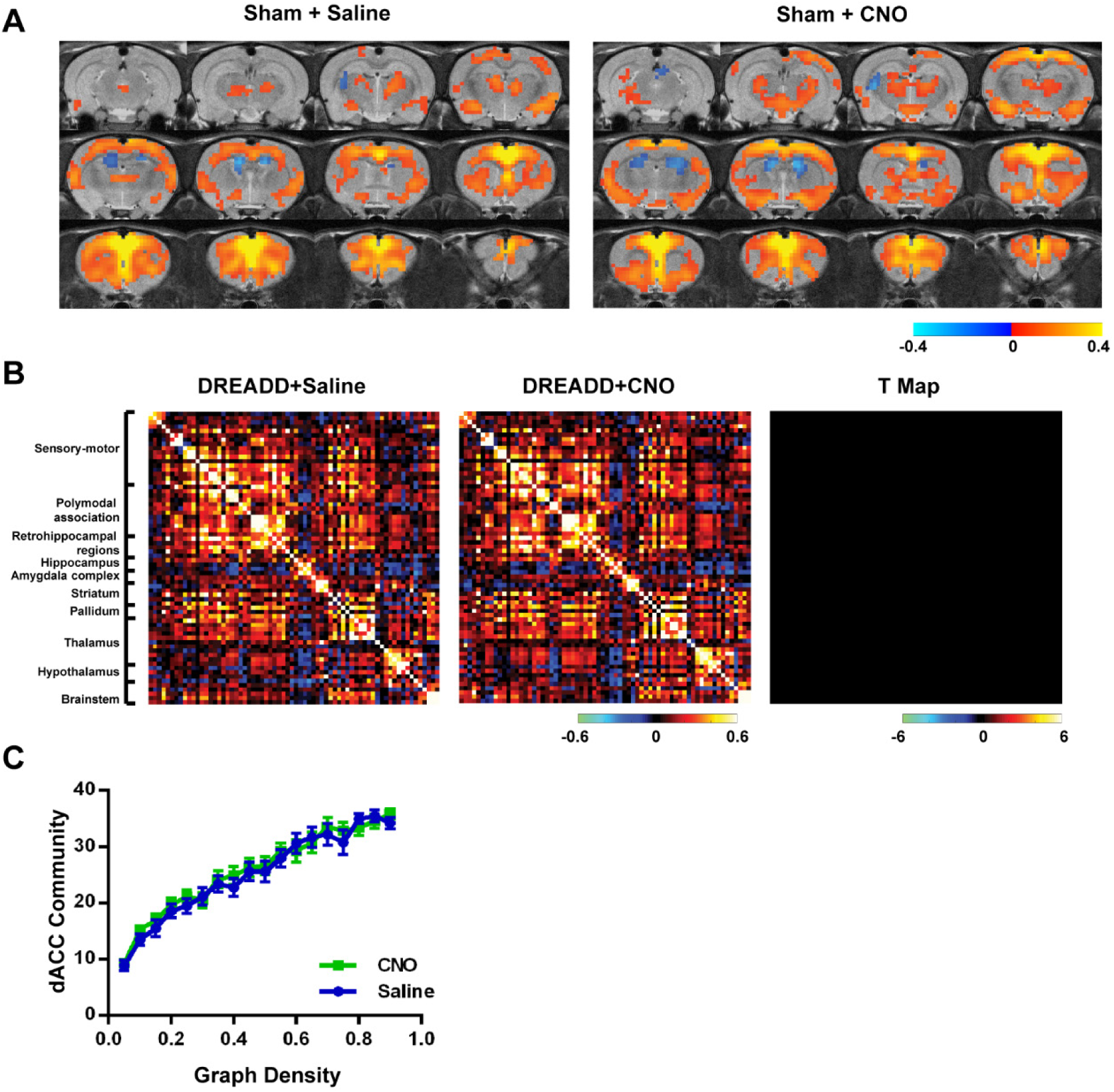
Sham rats did not show reduced RSFC or dACC community size. **A**) Averaged seedmaps after CNO and saline injections in the sham group (one sample t test, linear mixed model, p<0.0005, FDR corrected; n=8, 41scans); **B**) Averaged ROI-wise RSFC matrices after saline and CNO injections in the sham group. No connections exhibit RSFC changes (two-sample t-tests, linear mixed model, p < 0.05, FDR corrected). **C**) The size of the dACC-associated community as a function graph density after saline and CNO injections in the sham group (two-sample t-tests, linear mixed model, p > 0.05 for all densities tested).

**Figure S5.**
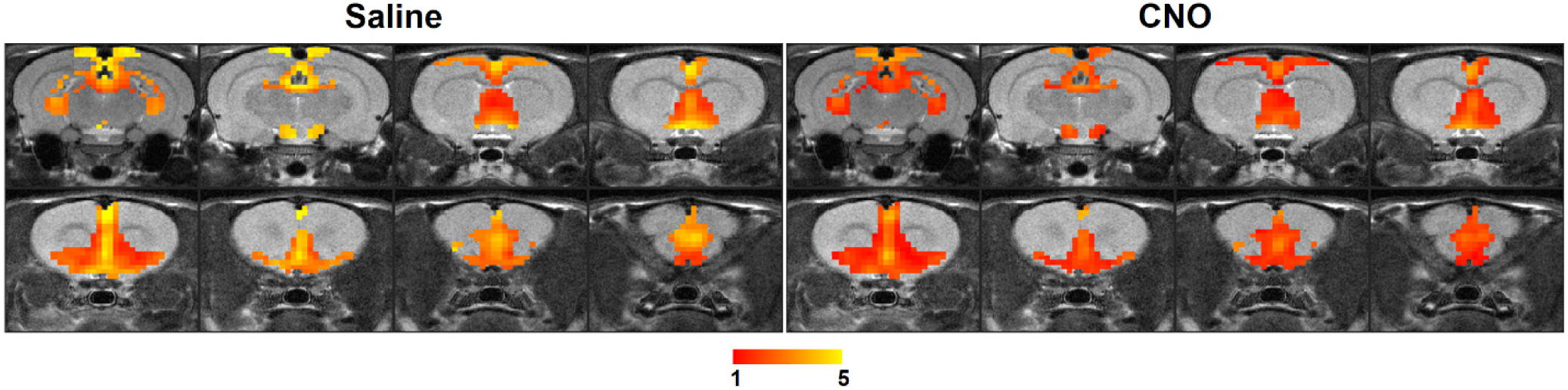
DMN pattern revealed by independent component analysis. rsfMRI data collected after saline and CNO injections were pooled together, and Group ICA was conducted. The DMN component was back projected to each group after group ICA.

**Figure S6.**
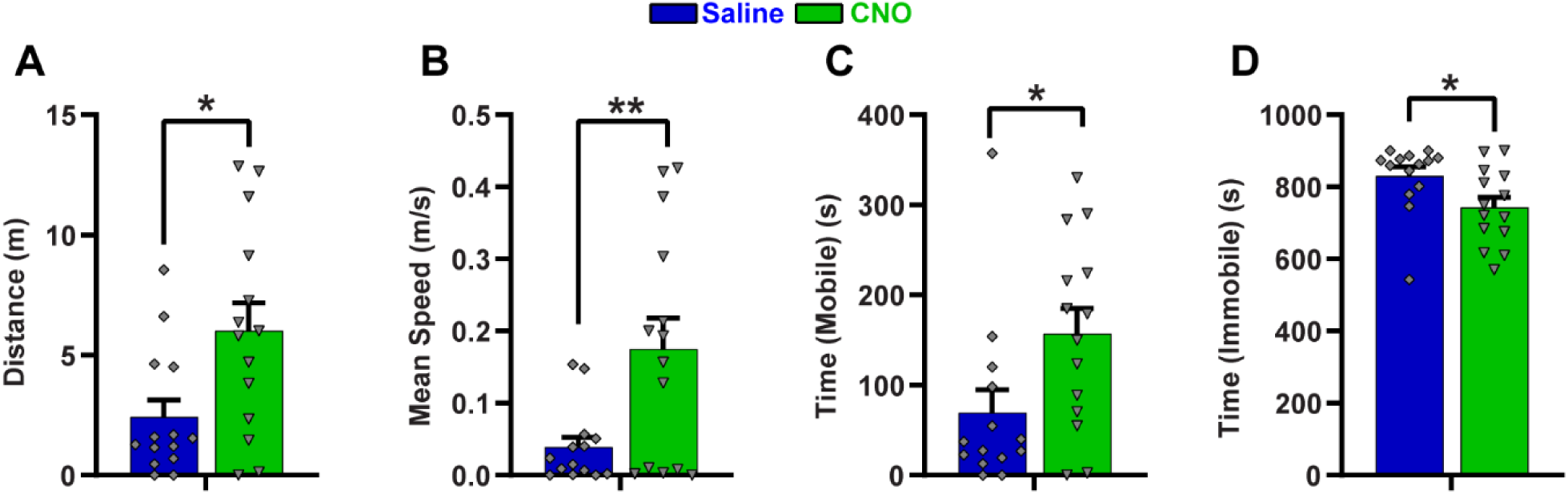
Suppressing dACC changed DMN-related behavior (n = 14). **A**) Total distance traveled; **B**) Mean speed; **C**) Total mobile time; and **D**) Total time when the animal was immobile for at least 2s in home-cage in the last 15 min of tests (i.e. 30 min after injection of CNO or Saline). *: p < 0.05, **: p < 0.01.

**Figure S7.**
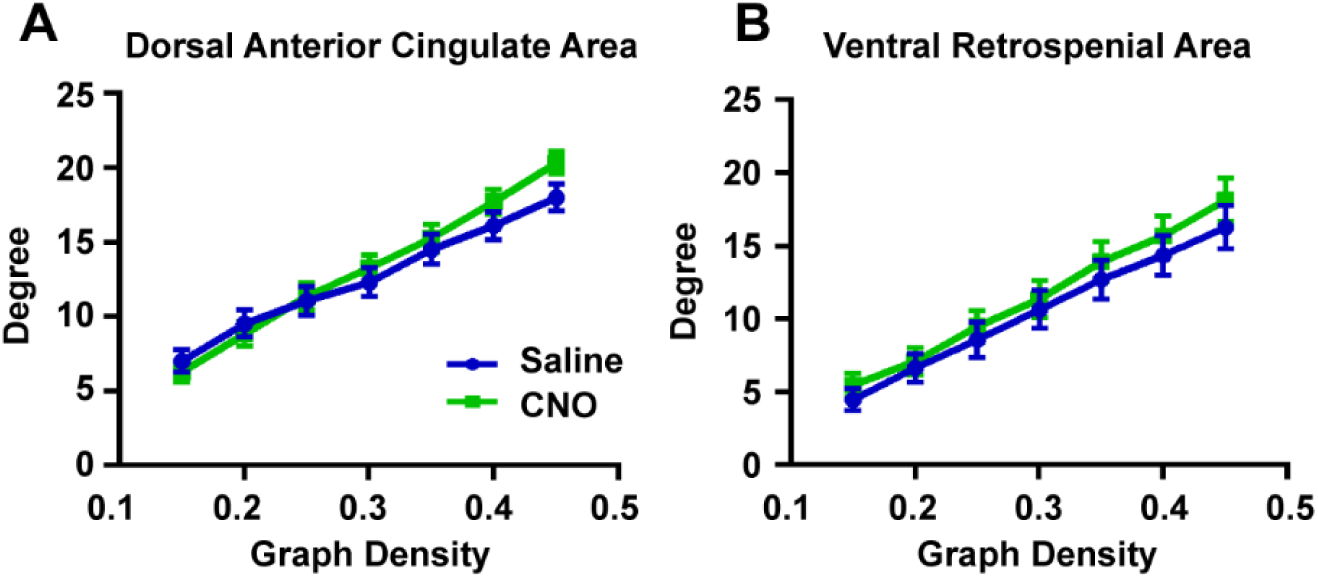
Sham group did not show changed degree of dACC or RSC. Degree changes of the dACC and ventral RSC in the sham group (two-sample t-tests, linear mixed model, p > 0.05 for all densities tested, n=8, 41 scans).

**Figure S8.**
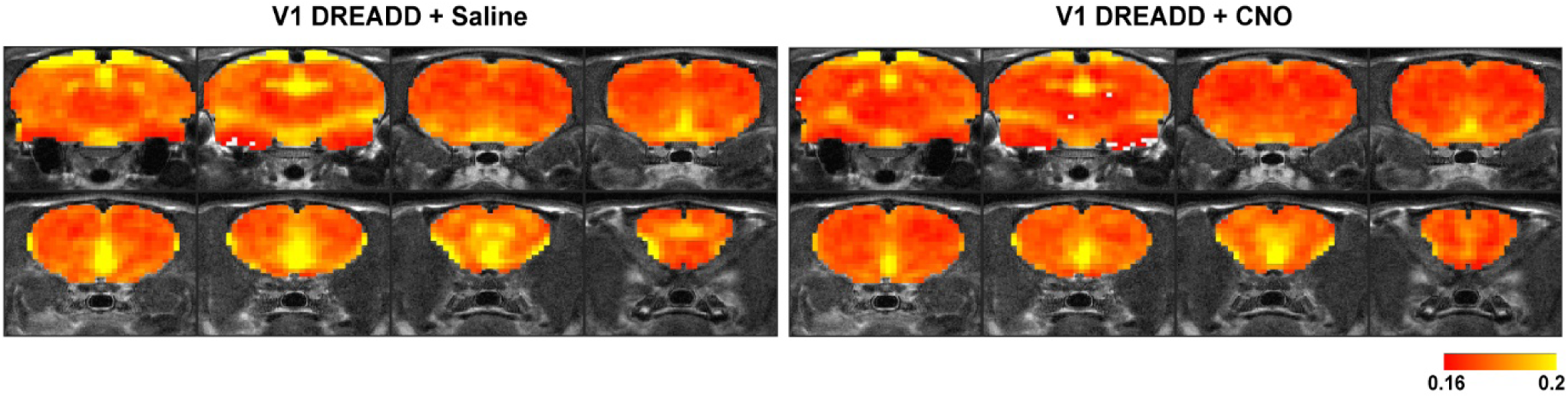
Suppressing V1 did not affect DMN activity. DMN constructed using fALFF in the V1 DREADD group after saline and CNO injections (n=8, 66 scans).

**Figure S9.**
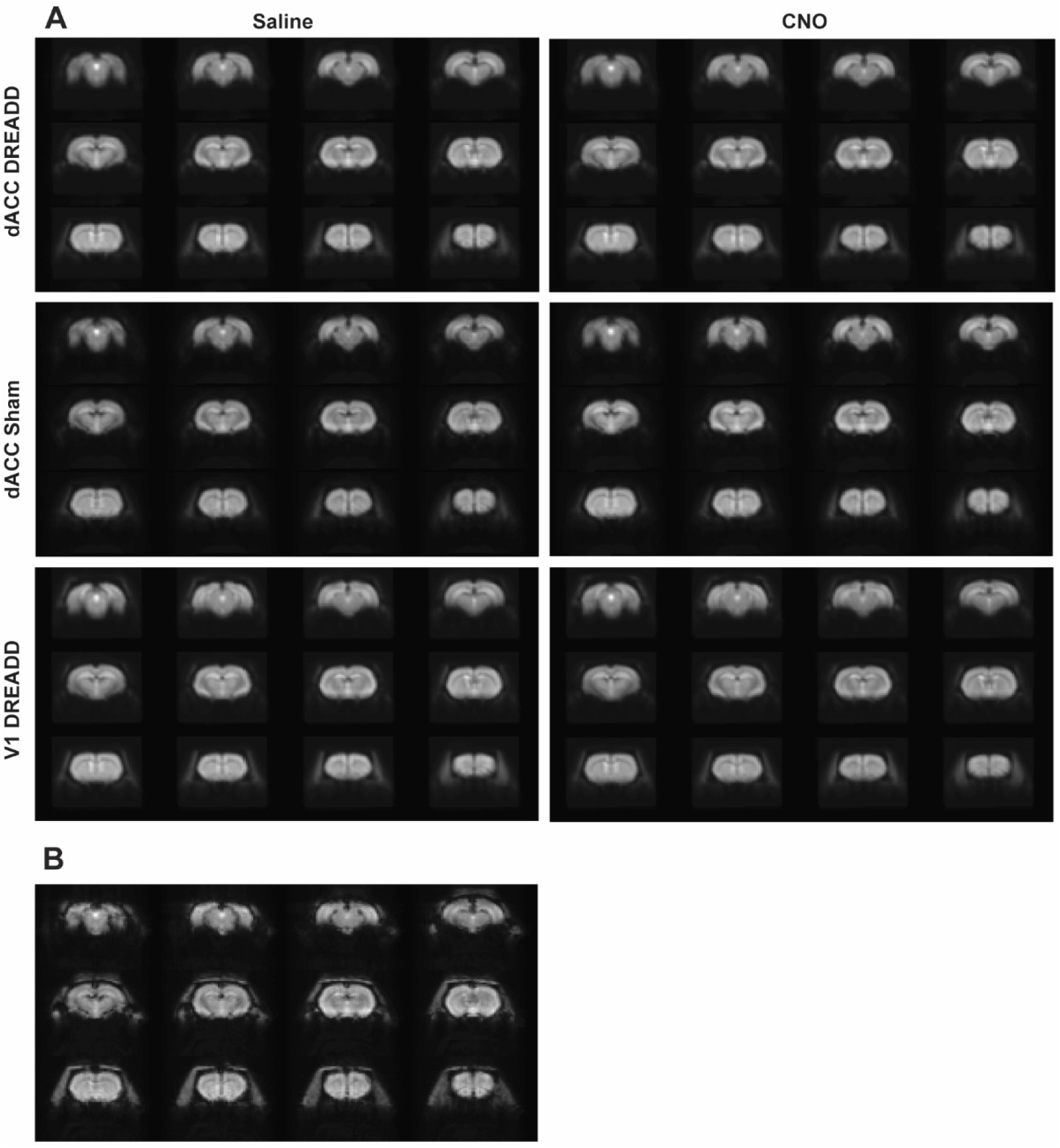
Raw EPI data. A. Averaged EPI image for each group and condition (dACC DREADD; dACC Sham; V1 DREADD); B. Example of a single EPI volume.

